# A pooling-based approach to mapping genetic variants associated with DNA methylation

**DOI:** 10.1101/013649

**Authors:** Irene M. Kaplow, Julia L. MacIsaac, Sarah M. Mah, Lisa M. McEwen, Michael S. Kobor, Hunter B. Fraser

## Abstract

DNA methylation is an epigenetic modification that plays a key role in gene regulation. Previous studies have investigated its genetic basis by mapping genetic variants that are associated with DNA methylation at specific sites, but these have been limited to microarrays that cover less than 2% of the genome and cannot account for allele-specific methylation (ASM). Other studies have performed whole-genome bisulfite sequencing on a few individuals, but these lack statistical power to identify variants associated with DNA methylation. We present a novel approach in which bisulfite-treated DNA from many individuals is sequenced together in a single pool, resulting in a truly genome-wide map of DNA methylation. Compared to methods that do not account for ASM, our approach increases statistical power to detect associations while sharply reducing cost, effort, and experimental variability. As a proof of concept, we generated deep sequencing data from a pool of 60 human cell lines; we evaluated almost twice as many CpGs as the largest microarray studies and identified over 2,000 genetic variants associated with DNA methylation. We found that these variants are highly enriched for associations with chromatin accessibility and CTCF binding but are less likely to be associated with traits indirectly linked to DNA, such as gene expression and disease phenotypes. In summary, our approach allows genome-wide mapping of genetic variants associated with DNA methylation in any tissue of any species, without the need for individual-level genotype or methylation data.

## INTRODUCTION

DNA methylation is an epigenetic mark that usually occurs at cytosine bases within CG dinucleotides (CpGs) in the human genome. CpGs often occur in dense clusters, known as CpG islands, which are surrounded by regions known as CpG shores; in non-malignant cells, CpG islands and shores are less frequently methylated than CpGs outside of these regions (Smith and Meissner 2013). Methylated CpG islands and shores in promoters may provide a “locking mechanism” that prevents repressed genes from being re-activated, while methylation in gene bodies is often associated with active transcription (Jones 2012; Wolf et al. 1984). In addition, DNA methylation has been linked to a wide range of diseases, including cancer, Alzheimer’s disease, bipolar disorder, and type 2 diabetes (Ambrosone et al. 2014; Baylin et al. 1998; Pease et al. 2013; The Cancer Genome Atlas Network, 2012; Baylin and Herman 2000; Irier and Jin 2012; De Jager et al. 2014; Lunnon et al. 2014; Gamazon et al. 2012; Dayeh et al. 2013). Interestingly, CpGs whose methylation has been associated with gene expression and disease are found not only in promoter regions or gene bodies but also in other parts of the genome, such as enhancers and insulators, suggesting additional roles of DNA methylation in transcriptional regulation (Jones 2012; Gutierrez-Arcelus et al. 2013; Zhang et al. 2014; Banovich et al. 2014; You et al. 2011). Although many studies have investigated potential additional roles, general conclusions about the role of methylation outside of promoters and gene bodies are still lacking (Jones 2012).

Several studies have investigated the relationship between DNA methylation and other epigenetic factors, such as histone modifications and chromatin accessibility (Wagner et al. 2014; Wrzodek et al. 2012; Shi et al. 2014; Zhang et al. 2014). For example, DNA methylation is associated with transcription factor (TF) binding (You et al. 2011; Thomson et al. 2010; Shi et al. 2014; Feldmann et al. 2013; Heyn 2014; Ziller et al. 2013; Wiench et al. 2011; Smith et al. 2014). In associations with epigenetic variation, the direction of causality is usually unclear; DNA methylation may affect TF-binding or may be affected by it, or both may be determined by another factor (or any combination of these).

In addition to epigenetic variation, DNA methylation can also be associated with genetic variation (Gibbs et al. 2010; Fraser et al. 2012; Smith et al. 2014; Drong et al. 2013; Lam et al. 2012; Grundberg et al. 2013; Bell et al. 2012, 2011; Ambrosone et al. 2014; Wagner et al. 2014; Liu et al. 2013; Shi et al. 2014; Moen et al. 2013; Heyn et al. 2013; Bibikova et al. 2011; Gutierrez-Arcelus et al. 2013; Zhang et al. 2014; Zhi et al. 2013; De Jager et al. 2014; Lunnon et al. 2014; Banovich et al. 2014). Associations with genetic variants such as SNPs are qualitatively different from epigenetic associations with disease or gene expression because the causality is clear: Mendelian randomization ensures that an individual’s genotype is a random combination of parental alleles, and thus any associations must be due to the effects of genotype (in a study free of confounding factors) (Mokry et al. 2014). Genetic variants showing these associations are known as DNA methylation quantitative trait loci (mQTLs).

Nearly all published human mQTL studies have relied upon commercially available microarrays that interrogate either ~28,000 CpGs or ~480,000 CpGs, corresponding to 0.1% and 1.7% of CpGs in the genome, respectively (Gibbs et al. 2010; Fraser et al. 2012; Smith et al. 2014; Drong et al. 2013; Grundberg et al. 2013; Bell et al. 2012, 2011; Ambrosone et al. 2014; Wagner et al. 2014; Liu et al. 2013; Shi et al. 2014; Moen et al. 2013; Heyn et al. 2013; Gutierrez-Arcelus et al. 2013; Zhang et al. 2014; Zhi et al. 2013; Banovich et al. 2014). Not surprisingly, the number of CpGs associated with mQTLs is generally greater in the studies using the larger arrays. However, the remaining 98.3% of CpG sites, including nearly all CpGs outside of CpG islands and shores, have yet to be included in any mQTL study, thus limiting our understanding of the roles of DNA methylation far from CpG islands.

In addition to covering only a small subset of the genome, DNA methylation arrays can be affected by experimental variability between samples, such as differences between the quality of bisulfite treatment across samples and “batch effects” that affect groups of samples, such as the day of hybridization (Leek et al. 2010; Sun et al. 2011). While many computational methods have been developed to mitigate these issues, they can be impossible to correct for perfectly and thus remain important potential sources of both false positives and false negatives (Stegle et al. 2012; Mostafavi et al. 2013; Johnson et al. 2007; Sun et al. 2011; Yousefi et al. 2013; Leek et al. 2010).

Another property of DNA methylation arrays is that they provide only the average methylation level for each targeted CpG in a sample. If an individual’s two alleles have the same methylation level, this is not an issue; however, samples heterozygous for a cis-acting mQTL will have allele-specific methylation (ASM) because cis-acting variants affect methylation of only the CpG allele to which they are linked (Kerkel et al. 2008; Shoemaker et al. 2010). Thus a significant source of information present in heterozygotes is lost. A previous ChIP-seq study used allele-specific information with a combined haplotype test, in which the authors modeled the read depth for each allele from each individual and tested whether there is a significant difference between allelic read depths across individuals, identifying thousands of histone modification QTLs (McVicker et al. 2013). Other ChIP-seq studies have also identified allele-specific histone modifications (Kasowski et al. 2013; Kilpinen et al. 2013; Mikkelsen et al. 2007), underscoring the value of allele-specific information.

Deep sequencing of bisulfite-converted DNA (Frommer et al. 1992) overcomes many of the limitations of microarrays by allowing genome-wide detection of DNA methylation and ASM (Lister et al. 2009; Xie et al. 2012; Lister et al. 2013; Schmitz et al. 2013a, 2013b). Applying this technology to F1 hybrid mice (CAST × 129) revealed over 10^5^ sites of allele-specific DNA methylation, suggesting that cis-acting effects of genetic variation on DNA methylation are widespread (Xie et al. 2012). However, performing whole-genome bisulfite sequencing — or even reduced representation bisulfite sequencing, which preferentially targets CpG-rich regions (Meissner et al. 2005, 2008) — on more than a few human individuals is prohibitively expensive. Nevertheless, because distal non-coding variants affecting DNA methylation may play key roles in some human traits (Ward and Kellis 2012; Dayeh et al. 2013; Grundberg et al. 2013), whole-genome bisulfite sequencing of many individuals may prove to be essential in understanding the genetic and epigenetic bases of phenotypic variation.

In this study, we developed a novel method for identifying mQTLs from pooled sequencing data that allows genome-wide mQTL mapping while reducing the effort, cost, and experimental variability associated with bisulfite sequencing and genotyping of many individual samples. Our approach allows us to study thousands of molecular traits in a single experiment, in contrast to existing pooled QTL mapping methods that are limited to individual phenotypes (Ehrenreich et al. 2010; Michelmore and Paran 1991). We first tested our approach in simulations and then applied it to bisulfite sequencing data from a pool of 60 human lymphoblastoid cell lines (LCLs). We identified over 2,000 novel mQTLs, including some that are also associated with variation in DNase hypersensitivity, TF-binding, gene expression levels, or complex diseases (Pickrell et al. 2010; Lappalainen et al. 2013; Degner et al. 2012; Welter et al. 2014; Ding et al. 2014). We also show that TF-binding sites and open chromatin regions from LCLs are enriched for mQTLs. Our approach represents a powerful and cost-effective framework for mapping mQTLs genome-wide, in any species.

## RESULTS

### A pooling approach to mQTL mapping

In most genetic association studies of quantitative traits, each sample is genotyped at many variants (typically at least 10^6^ for humans), and trait values (e.g. DNA methylation at specific CpGs) are measured for each sample. Genotypes are then compared to the trait values to detect statistically significant associations. This approach has been widely used in many species to map loci associated with both molecular-level traits (expression QTLs, mQTLs, DNase hypersensitivity QTLs, etc.) and organismal-level traits (height, blood pressure, etc.).

Our pooling approach is outlined in Figure 1. The central idea is that, for any cis-acting mQTL-CpG pair, if both the variant and the CpG are observed on the same DNA sequencing read, then any association between the two can be detected using Fisher’s exact test (see Methods). For example, a CpG near a G/T SNP might have a higher methylation level when linked to the G allele than when linked to the T allele. Analysis at the level of alleles, rather than individuals, allows samples to be pooled prior to bisulfite treatment because the allelic identity of every informative read can be inferred directly from the read’s sequence. In addition to minimizing experimental variability between samples, pooling obviates the needs for individual-level genotyping and DNA methylation profiling.

**Figure 1:**
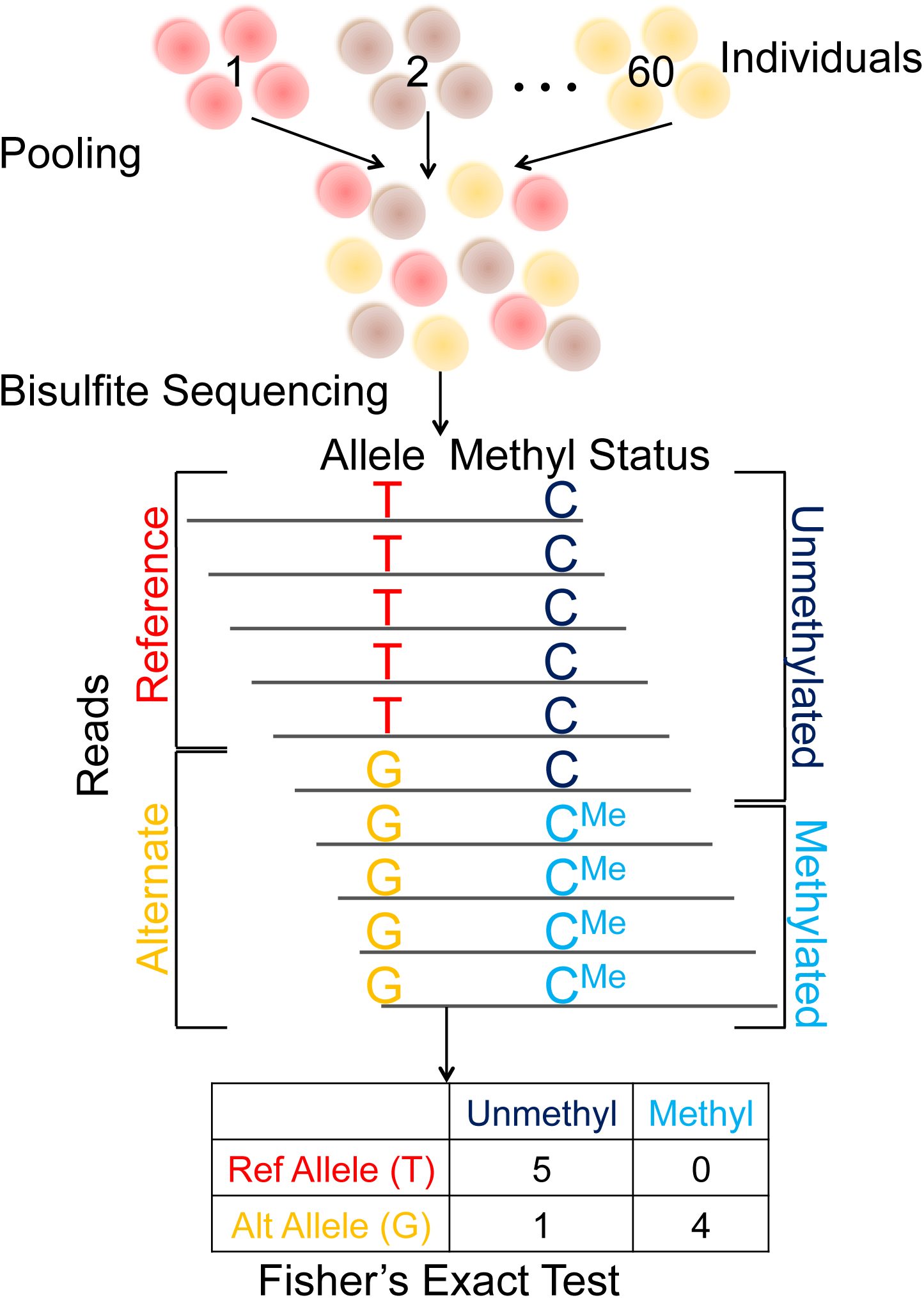
Using pooled bisulfite sequencing to identify mQTLs. Our pooling method enables us to identify mQTLs directly from bisulfite sequencing reads. In this approach, cells or DNA samples from all individuals are combined into a single pool, which is then subject to bisulfite sequencing. Alleles and methylation statuses are inferred from the sequence reads, which are then used to generate a 2×2 contingency table (where columns represent methylation statuses and rows represent alleles). Fisher’s exact test is used to compute a p-value for the null hypothesis of no association.

### Assessing the pooling approach via simulations

We compared our pooling approach (“pooled ASM method”), which allows us to detect allele-specific methylation (ASM), to a more traditional approach (“traditional non-ASM method”), which consists of bisulfite sequencing and genotyping of each sample separately, followed by the comparison of the individual-level genotypes and average DNA methylation levels. We focused on genome-wide methods and therefore did not directly include DNA methylation arrays in the simulations. Each simulation featured an mQTL-CpG pair, for which we estimated the power to detect the mQTL (at p < 0.001). Across simulations we varied five parameters: the effect size of the mQTL (the correlation between allele and methylation status), the read depth at the site (across all individuals), the minor allele frequency (MAF), the minor DNA methylation status frequency (the frequency of the less common methylation status), and the number of individuals. We also ran our simulations at additional p-value cutoffs and simulated variant-CpG pairs with no association between allele and methylation status at various p-value cutoffs to evaluate our false positive rate.

The pooled ASM method identified substantially more mQTLs than the traditional non-ASM method at most parameter values — the exception being a strong mQTL covered by many reads, in which case the methods both had close to 100% power (Figure 2, Supplemental Figures 1-8). In fact, for effect sizes ≤ 0.83, read depths ≤ 40x, and ≤ 100 individuals, the pooled ASM method had over twice the power of the traditional non-ASM method. The false positive rate for both methods was extremely low, especially for lower p-value cutoffs (Supplemental Figures 9-12). In addition, for nearly all parameter settings, the pooled ASM method produced significantly lower mQTL p-values than the traditional method (Supplemental Figure 13). We found similar results using two alternative statistical approaches to analyze the simulated data (Supplemental Figures 3-4).

**Figure 2:**
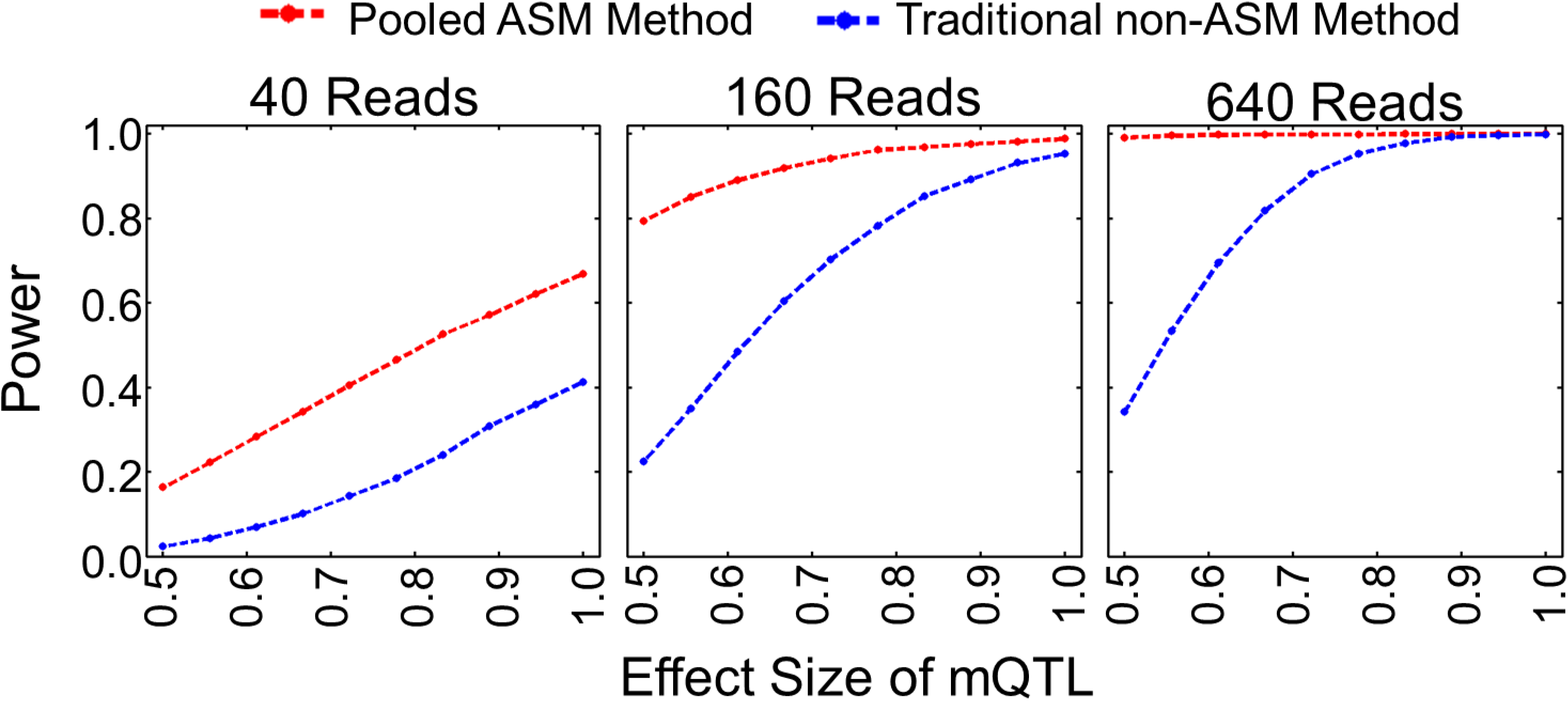
Simulations comparing our pooled ASM method to the traditional non-ASM method. Our pooled ASM method has more power to detect mQTLs than the traditional non-ASM method, especially for low effect sizes. Simulated reads from 100 individuals were sampled from a negative binomial distribution, with minor allele and minor methylation frequencies of 0.1. Power to detect an mQTL is shown for each method as a function of mQTL effect size and total read depth. The effect size is the correlation between allele and methylation status, and the power is the fraction of the simulations in which we identified an mQTL (with p < 0.001). Additional simulations are shown in Supplemental Figures 1-8, and ROC curves from the simulations are shown in Supplemental Figures 9-12.

The increased power of our approach is due primarily to the difference in what information is extracted from heterozygotes. For example, consider a strong mQTL (0% methylation of the major allele, 100% for the minor allele) with a low MAF that is present as only either major-allele homozygotes or heterozygotes in a particular cohort. The traditional non-ASM method averages together the effects of the two different alleles in each heterozygote, thus diluting the signal from each allele (the traditional method cannot distinguish between when the minor allele in heterozygotes always corresponds to the minor methylation status and when the allele in heterozygotes has no correlation with the methylation status). As a result, the two quantities being compared — average methylation of the CpG in homozygotes vs. heterozygotes — would be 0% vs. 50%. In contrast, because the pooled ASM method takes into account the allelic origin of each read, it is equally informative regardless of whether a read is from a homozygote or heterozygote. In our example, the two quantities in the comparison — methylation of the CpG in one allele vs the other — would be 0% vs. 100% in the pooled ASM method, which is more easily detected with limited data. This concept applies to any mQTL that can be found in a heterozygous state, even when all three possible genotypes are present in a cohort. As a result, the pooled ASM method had the greatest advantage at large MAFs (Supplemental Figures 5-6, Supplemental Figures 11-12) because these have the most heterozygotes.

### Applying our pooling approach to empirical data

As an experimental test of our pooling approach, we performed bisulfite sequencing on pooled genomic DNA from LCLs derived from 60 Yoruban (YRI) HapMap individuals (see Methods) (Frazer et al. 2007). We obtained ~860 million pairs of 101 bp reads, of which 77.1% passed our quality control filters and mapped uniquely, corresponding to an average per-base coverage of ~40x.

To assess the efficiency of our bisulfite treatment and pooling, we performed three quality control tests. First, to evaluate the efficiency of bisulfite conversion of unmethylated cytosines to thymines, we spiked unmethylated lambda phage DNA into the pool prior to bisulfite treatment. The percentage of unconverted lambda phage cytosines ranged from 0.0% to 0.2% across library/sequencing lane pairs, suggesting that the bisulfite treatment consistently had at least 99.8% efficiency, which is comparable to previous bisulfite sequencing studies (Banovich et al. 2014; Lister et al. 2009). Second, we estimated the rate of spurious cytosine to thymine conversion by creating two untreated control libraries from our pooled human samples. In these libraries, only 0.4-0.5% of bases that mapped to cytosines were thymines, suggesting a low rate of spurious C-to-T conversion, sequencing errors, and unannotated C/T SNPs. Finally, we estimated the relative abundance of each individual sample’s DNA in the pool and found that it was near the expected 1/60 ratio (1.0 – 2.8% abundance of each sample), confirming that our pool was relatively homogenous (Supplemental Figure 14). We note that sample heterogeneity should not impact our ability to map mQTLs so long as each allele is well-represented because our approach does not require an equal abundance of each sample in the pool.

Our pooled data provided sufficient coverage to test the strength of association at 823,726 variant-CpG pairs, from which we identified 2,379 mQTLs (at 2,332 unique CpGs) at p < 0.001 [corresponding to a false discovery rate (FDR) of 6.0%; see Methods and Supplemental Table 6]. Examples of two mQTLs are shown in Figure 3a. mQTLs tended to be close to their corresponding CpGs because our method required that they be on the same sequenced fragment; the average distance was 25.4 bp, with a range of 0-377 bp (Supplemental Figure 15).

**Figure 3:**
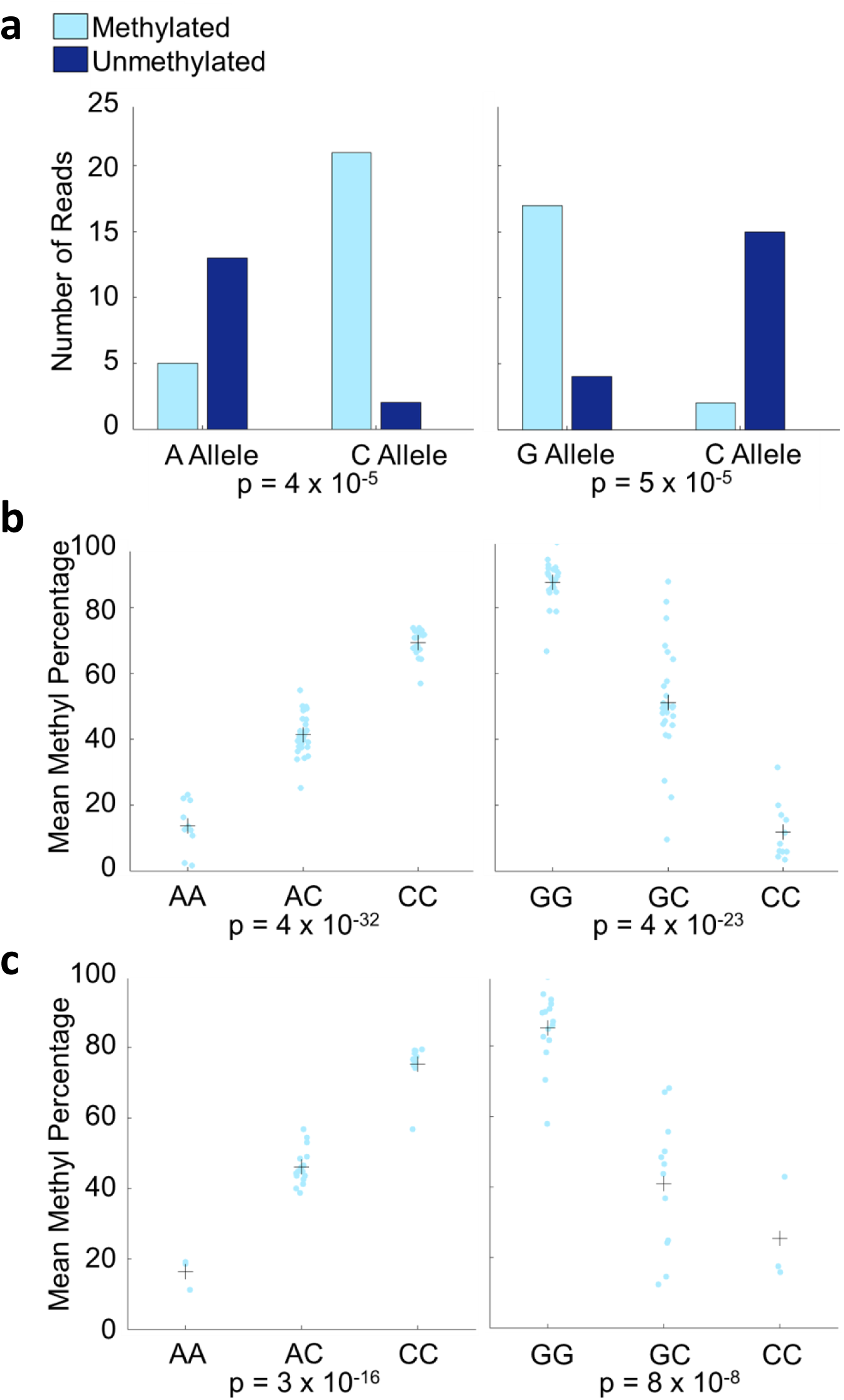
Pyrosequencing validation of mQTLs. Shown are two mQTLs involving SNPs previously identified in GWAS. (Left: The SNP is associated with age-related macular degeneration. Right: The SNP is associated with the ratio of visceral adipose tissue to subcutaneous adipose tissue.) **a)** Pooled bisulfite sequencing for the two mQTLs, showing strong association. **b)** Pyrosequencing validation of the two mQTLs in individual samples that were used for the pooled bisulfite sequencing confirms the bisulfite sequencing results. Light blue points are the methylation percentages from individuals, and crosses are the mean methylation percentages for individuals of each genotype. **c)** Pyrosequencing validation of the two mQTLs in 30 additional YRI individuals shows that the mQTLs are not limited to the individuals in our study. Light blue points are the methylation percentages from individuals, and crosses are the mean methylation percentages for individuals of each genotype.

We employed pyrosequencing on individual samples to validate our results in two ways. First, as a technical validation of the pooling method, we performed pyrosequencing at two CpGs with mQTLs in the same 60 samples that composed our pool. Both of these were successfully validated in the unpooled samples (Figure 3b). Second, we tested the reproducibility of our mQTLs by pyrosequencing eight CpGs (spanning a wide range of p-values and including the two with technical validation) in a separate set of 30 YRI samples, which are offspring of the 60 individuals in our pool. In seven out of eight cases, the mQTL was successfully validated in these additional samples (Figure 3c, Table 1, and Supplemental Figures 16-21; also described below).

**Table 1:** Pyrosequencing validation of mQTLs. Seven out of tested eight mQTLs were successfully validated using pyrosequencing. All mQTLs have p-values for the pyrosequencing validation on 30 YRI individuals who were not included in our pooled bisulfite sequencing data. mQTLs with two p-values are those that were also validated using the 60 YRI individuals in our study; the p-value from pyrosequencing of the 60 individuals in our data is listed first. All pyrosequencing data p-values have been Bonferroni-corrected.

| mQTL rsID | CpG position | mQTL annotation | p-value in pooled bisulfite sequencing data | p-value in pyrosequencing data |
| --- | --- | --- | --- | --- |
| rs10737680 | chr1:196679457 | GWAS hit | $4.3 \times 10^{-5}$ | $3.6 \times 10^{-32}$, $3.3 \times 10^{-16}$ |
| rs9517668 | chr13:99923789 | In LD with GWAS hit | $7.7 \times 10^{-4}$ | > 1 |
| rs12905925 | chr15:65164853 | dsQTL | $5.3 \times 10^{-6}$ | $4.0 \times 10^{-7}$ |
| rs617201 | chr17:3089145 | In LD with GWAS hit | $3.5 \times 10^{-5}$ | $6.4 \times 10^{-9}$ |
| rs1113144 | chr18:71774787 | In LD with GWAS hit | $1.4 \times 10^{-5}$ | $4.4 \times 10^{-13}$ |
| rs4809456 | chr20:61660870 | In LD with GWAS hit | $1.2 \times 10^{-4}$ | $2.4 \times 10^{-5}$ |
| rs62223713 | chr21:30158046 | Exon-level eQTL | $2.0 \times 10^{-4}$ | $9.7 \times 10^{-12}$ |
| rs7705033 | chr5:122774795 | GWAS hit | $4.7 \times 10^{-5}$ | $3.8 \times 10^{-23}$, $8.3 \times 10^{-8}$ |

We then compared our mQTLs to those in two recent studies of YRI LCLs (Zhang et al. 2014, Banovich et al. 2014), which both used the largest commercially available DNA methylation array, the Illumina Infinium HumanMethylation450 BeadChip (Bibikova et al. 2011). Comparing these studies to one another, we found 19.1-45.7% overlap (depending on the direction of analysis; see Methods). The disagreement was likely caused by a combination of false positives and false negatives (e.g. due to low power or between-sample variability). Comparing these two catalogs to our data, we found that less than 1% of our ~800K tested CpG sites were also tested in each of these studies. Focusing on the small number of sites in common, we found a similar level of overlap (40.0-53.3%; 4/10 and 8/15, respectively), which was significantly more than expected by chance (hypergeometric p = 5.5 × 10^−3^ and 9.6 × 10^−7^, respectively). In sum, our pooling-based approach agrees with microarray studies as well as these studies agree with one another, despite the major methodological differences between the approaches (Supplemental Figures 22-23).

### mQTLs were associated with molecular-level and organismal-level traits

To explore the potential effects of our mQTLs, we examined their distribution in CpG islands and shores as well as in genomic regions with different chromatin states (see Methods). Unlike DNA methylation microarrays that are primarily targeted in and around CpG islands, only 2.0% of our tested CpGs were in islands and only 10.8% were within 2 kb of CpG islands (Wu et al. 2010; Irizarry et al. 2009). Instead, most (86.0%) were in repressed/inactive genomic regions, which usually lack CpG islands. As a result, the majority (78.1%) of the CpGs with mQTLs was also in repressed/inactive regions, though this is less than the 86% expected by chance, suggesting that variants in active genomic regions may be more likely to influence methylation than those in inactive regions. Chromatin states that are related to active transcription and active enhancers were enriched for CpGs with mQTLs (Table 2, Supplemental Figure 24), suggesting that some of these mQTLs may affect transcription or enhancer activity.

**Table 2:** Enrichment of CpGs with mQTLs in LCL chromatin states. Most CpGs with mQTLs occur in regions of the genome that are in the “Repressed PolyComb” or “Quiescent/Low” chromatin states, but this is still no more than expected by chance. All p-values are Bonferroni-corrected. States with enrichment p < 0.05 are in bold. The corresponding fold-enrichments are in Supplemental Figure 24.

| Chromatin state | Percentage of CpGs with mQTLs in state | Enrichment p-value |
| --- | --- | --- |
| **1. Active TSS** | 0.90% | **$3.0 \times 10^{-6}$** |
| **2. Flanking Active TSS** | 1.0% | **$7.4 \times 10^{-3}$** |
| **3. Strong Transcription** | 3.8% | **$1.6 \times 10^{-2}$** |
| **4. Weak Transcription** | 3.3% | **$1.2 \times 10^{-4}$** |
| **5. Acetylated Active Enhancer** | 1.5% | **$1.7 \times 10^{-3}$** |
| 6. Acetylated Active Enhancer (Genic) | 0.26% | > 1 |
| **7. Active Enhancer** | 2.8% | **$4.4 \times 10^{-5}$** |
| 8. Active Enhancer (Genic) | 0.86% | $7.8 \times 10^{-2}$ |
| 9. Weak Enhancer | 3.1% | $4.9 \times 10^{-1}$ |
| 10. Weak Enhancer (Genic) | 0.99% | $6.6 \times 10^{-1}$ |
| 11. Bivalent TSS | 0.30% | > 1 |
| 12. Bivalent Enhancer | 1.1% | > 1 |
| 13. Repressed PolyComb | 23% | > 1 |
| 14. CTCF Only | 1.9% | $7.6 \times 10^{-2}$ |
| 15. Quiescent/Low | 55% | > 1 |

To understand if our mQTLs may be involved in cell-type-specific open chromatin, we compared our mQTL catalog with 402 DNase hypersensitivity experiments from the ENCODE and Epigenomics Roadmap projects, which measured chromatin accessibility (Thurman et al. 2012; The ENCODE Project Consortium 2012; Bernstein et al. 2010). We found that our mQTLs were more strongly enriched in regions of open chromatin in LCLs than in open chromatin in other cell types (p = 1.4 × 10^−7^). 9.9% of mQTLs are in open chromatin sites in LCLs, but only 4.9% of tested variants are in open chromatin sites from LCLs (fold-enrichment = 2.0); for all other cell types, 48.8% of mQTLs are found in open chromatin 11 regions, and 46.7% of tested variants are found in open chromatin regions (fold-enrichment = 1.0) (Supplemental Figure 25). This suggests that variants may be more likely to affect DNA methylation in tissues where they are in accessible chromatin.

To further understand the relationship between transcriptional activity and mQTLs, we intersected our mQTLs with TF-binding and histone modification ChIP-seq peaks from GM12878, an LCL studied extensively by the ENCODE consortium (The ENCODE Project Consortium 2012; Gerstein et al. 2012). We found that mQTLs were enriched in the binding sites for 12 TFs in GM12878; 4.6% of mQTLs are in binding sites of at least one of these TFs, while only 1.6% of tested variants are in these binding sites (total fold-enrichment across 12 TFs = 2.9) (Supplemental Table 1, Supplemental Figure 26). The strongest of these enrichments is for binding sites of CTCF, which often acts as an insulator by blocking the spread of chromatin states. SNPs associated with CTCF binding were recently mapped in LCLs (Ding et al. 2014); 18 of these are also mQTLs (p = 2.0 × 10^−3^), and increased binding is usually associated with decreased methylation (binomial p = 7.7 × 10^−4^), suggesting that variants affecting CTCF binding can also impact DNA methylation, consistent with their known relationship (Feldmann et al. 2013).

We next investigated the relationship between our mQTLs and other molecular traits by comparing our results to expression QTLs (eQTLs) and DNase hypersensitivity QTLs (dsQTLs) mapped in the same YRI population (Degner et al. 2012; Lappalainen et al. 2013). We found a strong overlap with dsQTLs, 48 of which were also mQTLs (p = 1.2 × 10^−27^, fold-enrichment = 7.4); alleles associated with lower DNA methylation tended to be associated with open chromatin (binomial p = 5.5 × 10^−7^). For eQTLs, we found that 28 mQTLs are exon-level eQTLs and five are gene-level eQTLs; for exon-level eQTLs, this was significantly more than expected by chance (p = 1.5 × 10^−3^, fold-enrichment = 1.8). In addition, considering SNPs in strong LD with eQTLs and dsQTLs revealed additional overlaps (Supplemental Table 2, Supplemental Table 3). We validated one eQTL overlap and one dsQTL overlap using pyrosequencing in an additional 30 YRI individuals (Table 1, Supplemental Figure 17, Supplemental Figure 21). These overlaps were consistent with our expectation of a relationship between DNA methylation, transcription, and chromatin structure (Gutierrez-Arcelus et al. 2013; Jones et al. 2013).

We also tested to what extent our mQTLs may impact organismal-level traits. To investigate this, we performed a similar comparison of our mQTLs with variants implicated by genome-wide association studies (GWAS) for diseases and other traits (Welter et al. 2014). Nine mQTLs were previously identified in GWAS or are in perfect LD with GWAS hits, and an additional 13 mQTLs are in strong (*r*^2^ ≥ 0.8) LD with GWAS-implicated variants (Welter et al. 2014) (Supplemental Table 4), though this is not significantly more overlaps than expected by chance (p = 0.67 for GWAS hits and variants in perfect LD; p = 0.70 for GWAS hits and variants in strong LD). We validated two of the overlaps using pyrosequencing in the individuals in our bisulfite sequencing study (Figure 3b, Table 1) and six of the overlaps in 30 additional YRI individuals; all but one test was successful (Figure 3c, Table 1, Supplemental Figure 16, Supplemental Figures 18-20). All of the validated mQTLs are associated with CpGs that are not included on commercially available microarrays, demonstrating that our method can identify novel disease-associated mQTLs.

Across the different types of QTLs/GWAS that we compared to our mQTLs, we noticed an interesting trend: QTLs associated with the earliest steps of transcription (dsQTLs associated with chromatin accessibility) had the strongest overlap with mQTLs, while moving progressively towards phenotype, the enrichment decreased at each stage (Figure 4). This suggests a buffering or dilution of mQTL effects at transcriptional and post-transcriptional levels, consistent with recent models for the attenuated effects of eQTLs on protein levels (Battle et al. 2014). As such, this could allow for plasticity in DNA methylation, with a small subset of methylation changes overcoming the many layers of gene expression regulation to ultimately affect organismal phenotypes.

**Figure 4:**
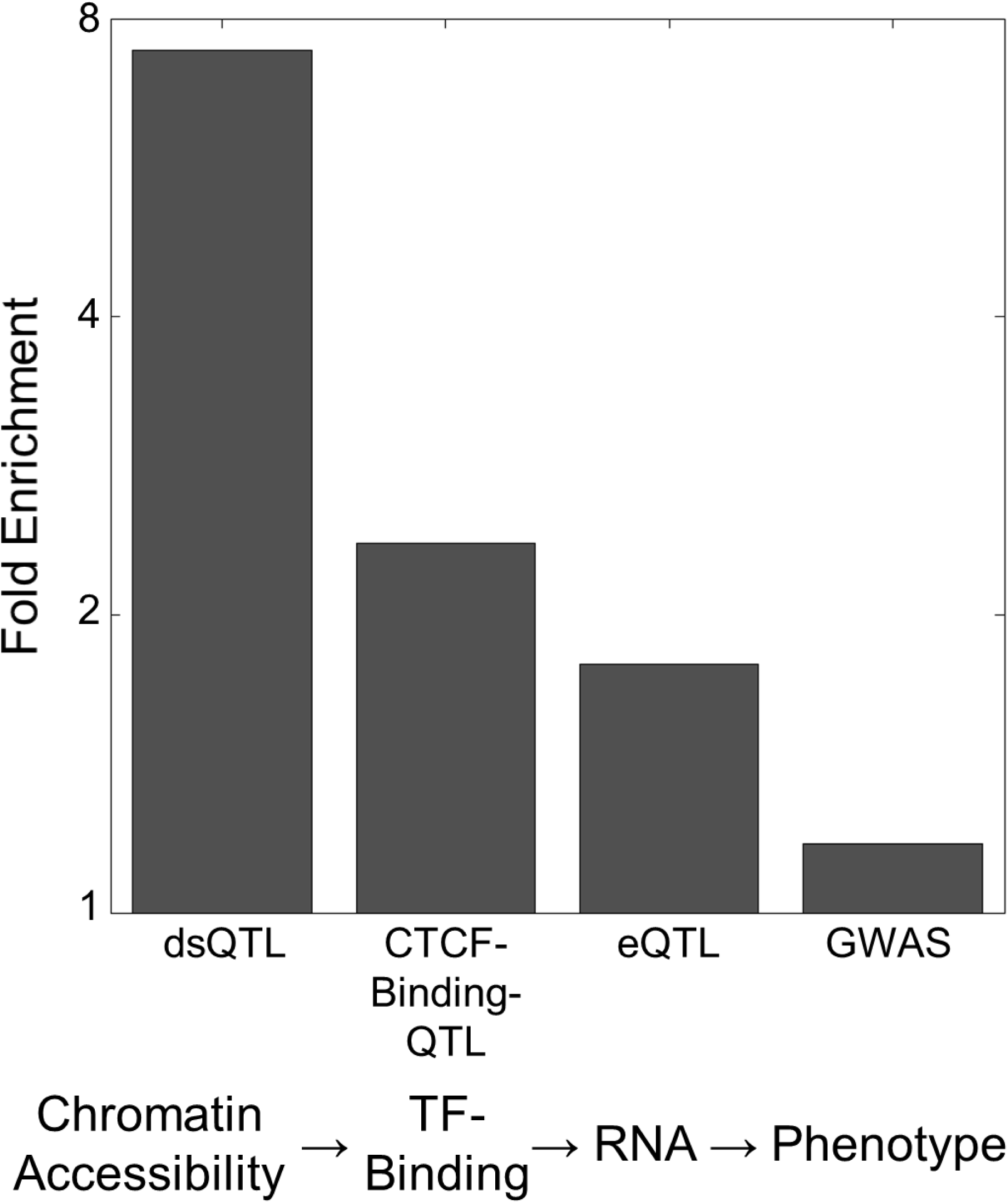
Fold-enrichments of mQTLs overlapping other QTLs and GWAS hits. mQTLs have greater enrichment for QTLs physically associated with DNA than they do for QTLs related to downstream traits only indirectly linked to DNA.

## DISCUSSION

The pooling approach introduced here represents a major advance in our ability to map cis-acting mQTLs genome-wide. Our simulations demonstrated the substantial increase in power that our approach provides as a result of accounting for ASM. In an empirical test of our approach, we identified mQTLs in many functionally important sites that are not covered by microarrays, including TF-binding sites, dsQTLs, CTCF-binding-QTLs, eQTLs, and GWAS hits. Thus, our approach overcomes two of the greatest weaknesses of microarrays — inability to detect ASM and limited coverage of CpGs — enabling us to detect thousands of novel mQTLs throughout the genome.

An additional advantage of our approach over microarrays and other individual-level assays is the minimization of between-sample experimental variability and batch effects, which are known to be important sources of error (Mostafavi et al. 2013). By pooling samples prior to bisulfite treatment, most potential sources of variability are eliminated. However, because our simulations did not account for these factors, we may have underestimated our increase in power over the traditional non-ASM method.

Several important advantages of the traditional approach of genotyping and using microarrays for DNA methylation measurement were also not captured by our simulations. For example, the traditional approach can detect mQTLs acting at any distance away from their CpG targets, including trans-acting mQTLs. Because pooling requires the variant and CpG to be on the same sequenced DNA fragment, the distance is limited by the size of these fragments. (This includes the insert size in paired-end reads, so the distance can be many kilobases if large fragments are selected for sequencing, though a large variance in insert size can lead to fewer reads covering a given variant/CpG pair.) A previous study reported that the majority of “likely causal” mQTL variants are within 100 bp of the corresponding CpG (Banovich et al. 2014); therefore, our approach should be able to detect most of these causal variants, though it will have less power to detect additional mQTL variants that are in LD with the causal variants, since these can be much further away. As read lengths increase with new sequencing technologies, this limitation may be lessened.

Another limitation of our approach is that it excludes C/T and G/A SNPs from consideration because their genotypes cannot be disentangled from the effects of bisulfite conversion. However, because most SNPs (including C/T and G/A SNPs) are in strong linkage disequilibrium with other variants, their mQTL associations may still be measured via other “tag SNPs” (that are on the same read as the corresponding CpG), as is done in any QTL or GWAS study. Moreover, this limitation is shared by other methods for inferring ASM from individual-level bisulfite sequencing data.

Our approach also cannot account for population structure or any known covariates (e.g. gender or environmental differences) since the individual from which each read originated is unknown. In our study, this is probably not an issue because our individuals lack significant population structure, and the cell lines were grown in controlled laboratory conditions. Future applications of this approach would ideally use cohorts in which these issues are minimized, such as unstructured natural populations or controlled F2 crosses (as is ideal for any QTL study or GWAS) (Bush and Moore 2012).

Finally, our approach relies on detecting alleles directly from reads as well as detecting the positions of variants in the genome directly from reads if the data are from populations for which variant positions are not known. Detecting alleles from reads is not perfect due to sequencing errors and imperfect mapping. However, this is unlikely to have a substantial effect on our results because only errors at the variant positions would lead to an incorrect allelic assignment. Detecting variants is a more challenging problem (Li 2014; Nielsen et al. 2011), but it can be minimized by limiting the genotype calls to known variant positions, by sequencing deeply, or by excluding rare variants.

While pooling individuals enables us to overcome limitations of previous approaches for mQTL mapping, we are not the first to use pooling for QTL mapping. Previous studies have used pooling for QTL or association mapping of individual traits. These approaches include bulk segregant analysis (BSA) (Michelmore and Paran 1991) and X-QTL mapping (Ehrenreich et al. 2010). They involve phenotyping many individuals for a specific trait followed by genotyping (via microarrays or sequencing) pools of individuals with either high or low trait values. QTLs are then identified as genetic variants with different allele frequencies between the two pools. These approaches have been gaining popularity because they, like ours, do not require individual-level genotyping, which is often the most laborious and expensive component of QTL mapping. In comparison to our method, BSA and X-QTL mapping can be applied to a much wider range of traits, including organismal-level traits. Two key advantages of our mQTL approach are that it can be applied to millions of molecular-level traits in one pool (as opposed to a single trait for every two pools) and that it does not require any individual-level phenotyping.

We are also not the first to leverage allele-specific information for QTL mapping. Previous work has also used allele-specific information in ChIP-seq data to map QTLs associated with histone modification levels (McVicker et al. 2013) and to identify allele-specific histone modifications (Kasowski et al. 2013; Kilpinen et al. 2013; Mikkelsen et al. 2007). Unlike ChIP-seq, bisulfite sequencing results in approximately even sampling of all DNA in the sample, not just the DNA that is bound by a protein of interest. Therefore, our goal is not to identify the difference in read depth between alleles but rather the difference in the fraction of CpG methylation between alleles. Because these problems require distinct approaches, we proposed a novel approach for use in bisulfite sequencing studies.

While our approach has many advantages over measuring genotypes and DNA methylation separately for each individual, there are ways to combine the approaches that would leverage ASM without sacrificing the ability to identify trans-acting and distal cis-acting mQTLs. For example, one could alter the step at which pooling takes place by creating a uniquely barcoded library for each sample prior to pooling and bisulfite treatment, thereby allowing each read to be assigned to its sample of origin. However, this would come at the cost of the additional experimental variability, effort, and expense associated with creating a separate library for each sample. Alternatively, even with no pooling, the extra power gained from allele-specific information in heterozygotes can be achieved by inferring both alleles and DNA methylation statuses directly from reads.

Large mQTL mapping studies have been limited by the effort and expense involved in data generation; our approach does not require individual-level data for either genotypes or DNA methylation, thereby significantly decreasing the barrier to mQTL mapping in any tissue or species. Although the number of mQTLs we identified was modest, this is primarily a limitation of our sequencing depth and not the method itself. As illustrated by our simulations, with 40x coverage, our power to detect even strong mQTLs is modest. As sequencing becomes less expensive, this pooling approach may help us achieve a comprehensive understanding of the relationship between genetic variation and DNA methylation, which will provide insight into traits such as evolutionary adaptations in many species and human diseases in many tissues. Moreover, with some modifications, the pooling framework we have introduced could also be applied to mapping QTLs for other molecular-level traits, such as TF-binding and histone modifications. We anticipate that pooling will enable us to leverage sequencing technology to study the relationship between genetic and epigenetic variation in a much wider range of cell types and species than has previously been possible.

## METHODS

### Simulations comparing the pooled and traditional association methods

In order to compare the power of identifying mQTLs using the pooled ASM method – sequencing on pooled samples and accounting for ASM – versus the traditional non-ASM method of associating genotypes with average methylation statuses, we performed simulations of a single theoretical variant-cytosine pair, varying several parameters of interest. We varied the effect size (correlation between the variant’s allele and the cytosine’s methylation status), read depth (number of reads across all individuals), MAF and minor methylation status frequency (the minor methylation status is the cytosine methylation status with fewer reads), and number of individuals. We tested effect sizes ranging from 0.5 to 1.0; 20, 40, 80, 160, 320, 640, and 1280 reads across individuals; minor allele and methylation status frequencies 0.1, 0.2, 0.3, 0.4, and 0.5; and 25, 50, 100, 200, and 400 individuals. When assigning genotypes to individuals, we always assumed that our population is in Hardy-Weinberg equilibrium. For each parameter combination, we ran 10,000 simulations for the variant-cytosine pair.

We ran each simulation as if we were randomly selecting DNA fragments from a pool of cells. For the pooled data simulations, we sampled allele-methylation status pairs with replacement for each read. We used Fisher’s exact test, implemented in Matlab using Michael Boedigheimer’s fexact function (www.mathworks.com/matlabcentral/fileexchange/22550-fisher-s-exact-test), to quantify the association. For the traditional non-ASM method simulations, we randomly assigned each read to an individual so that each individual had approximately (but not necessarily exactly) the same number of reads. For each individual, we used the genotype that we had initially assigned to it and computed the average methylation status, rounded to the nearest integer. We used a 2 × 3 Fisher’s exact test to compute the associations between genotype and average methylation status, which was implemented in Matlab using Giuseppe Cardillo’s myfisher23 function (www.mathworks.com/matlabcentral/fileexchange/15399-myfisher23). For comparison, we also computed p-values using the p-value of the F-statistic for the regression that predicts methylation status as a function of allele/genotype and using the asymptotic p-value of the Pearson correlation between allele and methylation status; when we ran these simulations for the “traditional non-ASM method,” we did not round the average the methylation status for each individual. For each method, we estimated power as the fraction of simulations in which the variant-cytosine pair reached p < 0.001, the same p-value threshold that we used for identifying mQTLs in our real data. We then ran additional simulations, as described in the Supplemental Methods.

### Whole-genome bisulfite sequencing library preparation

We pooled genomic DNA derived from 60 Yoruban LCLs (parental samples from 30 HapMap trios, purchased from Coriell). 48 μg of this DNA was spiked with 240 ng unmethylated cl857 Sam7 Lambda DNA (Promega, Madison, WI) to yield 0.5% W/W lambda DNA. The DNA was fragmented with a Covaris instrument (Covaris) in 50 uL volumes (duty factor 10%, peak incident power 175,200 cycles per burst, 40 seconds duration, 5.5-6.0°C), followed by end repair, adenylation, and adapter ligation using the TruSeq DNA LT Sample Prep Kit (Illumina) as per manufacturer’s instructions. Purification steps were performed using Agencourt Ampure beads (Beckman-Coulter). All 24 indexed methylated adapters from TruSeq DNA LT Sample Prep Kit Sets A and B (Illumina) were used to construct the libraries in order to increase base complexity.

Adapter-ligated DNA of 400-500 bp was isolated by 2% agarose gel electrophoresis using low range ultra agarose (Bio-Rad) with SYBR Gold Nucleic acid gel stain (Invitrogen), and fractions were purified using MinElute Gel Extraction Kit (Qiagen). Two of the 24 libraries were reserved and not bisulfite-converted for control purposes. Sodium bisulfite conversion was carried out on each of the 22 remaining libraries using the EpiTect Bisulfite Kit (Qiagen #59104) as per manufacturer’s 2006 instructions, except the reaction mix incubation cycle from the Whole-Genome Bisulfite Sequencing for Methylation Analysis was used (Illumina, Part # 15021861 Rev. A) and consisted of 95°C for 5 min., 60°C for 25 min., 95°C for 5 min., 60°C for 85 min., 95°C for 5 min., 60°C for 175 min., 3 cycles consisting of 95°C for 5 min., and 60°C for 180 min., ending with a 20°C hold. Bisulfite-converted products were purified using MinElute PCR Purification Kit (Qiagen).

Adapter-ligated, bisulfite-converted libraries were enriched using KAPA HiFi HotStart Uracil+ ReadyMix uracil-insensitive polymerase (D-mark Biosciences #KK2801). Thermocycler parameters for bisulfite-converted libraries consisted of 98°C for 45 sec.; four cycles of 98°C for 15 sec., 65°C for 30 sec., 72°C for 30 sec., ending with a 4°C hold. Adapter-ligated, non-bisulfite-converted control libraries were enriched using the reagents and protocol from the TruSeq DNA LT Sample Prep Kit (Illumina). Thermocycler parameters consisted of 98°C for 30 sec.; four cycles of 98°C for 10 sec., 60°C for 30 sec., 72°C for 30 sec., ending with a 4 °C hold. PCR reaction products were purified using Agencourt Ampure beads (Beckman-Coulter). Library validation was performed using the KAPA SYBR FAST Universal qPCR Library Quantification Kit (D-mark Biosciences, #KK4824) to measure the concentration of viable sequencing template molecules as well as the Agilent Bioanalyzer High Sensitivity DNA Assay (Agilent) to determine the size and distribution of the template molecules. Libraries were further concentrated using the MinElute PCR Purification Kit (Qiagen). Each non-control library was divided across eight sequencing lanes in three flowcells, and each control library was divided across four of these sequencing lanes, which were in two of the flowcells.

### Data processing

To obtain high-quality allele and methylation information from our bisulfite sequencing data, we trimmed reads, aligned reads to the genome, identified methylation statuses for cytosines and alleles for SNPs, and filtered our data according to various metrics. Details are described in the Supplemental Methods.

### Identifying methylation quantitative trait loci (mQTLs)

To find cytosines that are strongly associated with genetic variation, we combined variants in perfect linkage disequilibrium (LD), tested whether each association was significant, and used permutation tests to compute a false discovery rate (FDR). We associated every variant (including SNPs, insertions, and deletions) with every cytosine on the same read, creating a list of variant-cytosine pairs; thus, CpGs that never occurred on a read with a variant were not analyzed. When associating variants with CpGs, we combined cytosines on different strands of each CpG. For each cytosine, we combined associated variants whose genotypes in 1000 Genomes AFR (The 1000 Genomes Project Consortium, 2010) are perfectly correlated by replacing the allele of the variant that is later on the chromosome with the associated allele for the variant earlier on the chromosome (Supplemental Figure 28). After combining SNPs, we computed the significance of the associations between variants’ alleles and cytosines’ methylation statuses for every variant-cytosine pair using Fisher’s exact test (Fisher 1922), which we implemented using Scipy’s fisher_exact (Oliphant 2007). We removed all variant-cytosine pairs for which the number of reads with the minor allele or minor methylation status was so low that, given the total number of reads, the lowest p-value that could be achieved using Fisher’s exact test was ≥ 0.001. For each cytosine, we defined methylation quantitative trait loci (mQTLs) to be variants for which the association between the allele and the cytosine’s methylation status had a Fisher’s exact test p < 0.001; this gave us an FDR of 0.0601. We computed the FDR by permuting the methylation statuses for each variant-cytosine pair 100 times, re-computing the p-value for each pair, re-identifying mQTLs, and averaging the number of mQTLs across permutations. The FDR is the ratio of the average number of mQTLs across permutations to the number of real mQTLs. To check for possible position-specific biases in our mQTL mapping, we intersected the variant-cytosine pairs for our real mQTL list with the variant-cytosine pairs for each permuted mQTL list; no variant-cytosine pair for a real mQTL occurred in more than two permuted lists, as expected.

### Computing mQTL enrichments

In order to test whether mQTLs associate with specific chromatin states or transcription factor (TF)-binding sites, we intersected our CpGs with mQTLs with chromatin states from GM12878 (Kasowski et al. 2013; Ernst and Kellis 2012) and our mQTLs with reproducible (IDR 2%) TF-binding sites and histone modification regions from GM12878 (Myers et al. 2011; The ENCODE Project Consortium 2012; Gerstein et al. 2012; Landt et al. 2012; Li et al. 2011). If multiple data-sets were available for a TF or histone modification, we used the dataset with the largest number of peaks. We used the hypergeometric test, with the background defined as all variants in variant-cytosine pairs that were tested for being mQTLs. We corrected p-values using the Bonferroni correction (Shaffer 1995). For chromatin state enrichments, we multiplied p-values by the number of possible chromatin states. For the TF-binding site and histone modification enrichments in GM12878, we multiplied p-values by the number of TFs and histone modifications tested. In addition to computing p-values, we computed fold-enrichments for chromatin states, TF-binding sites, and histone modification regions.

To test whether mQTLs associate with open chromatin in various cell types, we intersected the variants we tested for mQTLs with Joseph Pickrell’s list of SNPs in open chromatin regions in various cell types, which were downloaded from https://github.com/joepickrell/1000-genomes (Bernstein et al. 2010; The ENCODE Project Consortium 2012; Thurman et al. 2012). We used a hypergeometric test to evaluate the enrichment for mQTLs in variants overlapping open chromatin regions in each cell type. We then used a one-sided Wilcoxon rank-sum test to compare the mQTL enrichments for LCLs versus other cell types, where our null hypothesis was that the median hypergeometric p-value for LCLs is not less than the median hypergeometric p-value for other cell types.

### Computing overlap between mQTLs, eQTLs, dsQTLs, CTCF-binding-QTLs, and GWAS hits

In order to evaluate whether mQTLs may be associated with gene expression, open chromatin, CTCF-binding, and organismal-level traits, we computed overlap between mQTLs and variants from expression QTLs (eQTLs), DNase hypersensitivity QTLs (dsQTLs), CTCF-binding-QTLs, and genome-wide association study (GWAS) datasets. To compute the enrichment of mQTLs in another molecular QTL study, we used a hypergeometric test, with the background defined as all variants tested in both our study and the other study. In addition to computing p-values, we computed fold-enrichments. The p-values and fold-enrichments reported in the Results section are those for molecular QTLs and variants in perfect LD with molecular QTLs.

Because we do not know all of the SNPs in our study that have been tested in a GWAS, we applied a method similar to the one used by Lappalainen *et al.* (Lappalainen et al. 2013) for finding the enrichment of mQTLs in GWAS hits. We first created 10,000 null sets of variants that were tested for being mQTLs in our study, where each variant *i* in a null set had an MAF within 0.001 of variant *i* in the set of mQTLs found in our study. Then we found the number of overlaps between the null set and GWAS hits. The p-value was defined as the fraction of null sets with at least as many overlaps with GWAS hits as mQTLs have with GWAS hits. We computed the fold-enrichment for GWAS in the same way as the fold-enrichment for molecular QTLs.

### Validation of mQTLs through bisulfite pyrosequencing

Bisulfite PCR-pyrosequencing assays were designed with PyroMark Assay Design 2.0 (Qiagen). The regions of interest were amplified by PCR using the HotstarTaq DNA polymerase kit (Qiagen) as follows: 95°C for 15 min. (to activate the Taq polymerase), 45 cycles of 95°C for 30 sec., 58°C for 30 sec., and 72°C for 30 sec., and a 72°C 5 min. extension step; primer sequences are listed in Supplemental Table 5. For pyrosequencing, a single-stranded DNA was prepared from the PCR product with the Pyromark™ Vacuum Prep Workstation (Qiagen), and the sequencing was performed using sequencing primers on a Pyromark™ Q96 MD pyrosequencer (Qiagen). The quantitative levels of methylation for each CpG dinucleotide were calculated with Pyro Q-CpG software (Qiagen). p-Values for associations were the asymptotic p-values of the correlations between genotype and average methylation from the pyrosequencing assay. We performed pyrosequencing on the 60 individuals in our pool as well as on 30 additional individuals who are the offspring of those 60.

## DATA ACCESS

The data generated in this study have been submitted to the NCBI Sequence Read Archive (SRA; http://www.ncbi.nlm.nih.gov/sra) under accession number SRP045408.

## ACKNOWLEDGMENTS

We would like to thank M. Hirst, M. Moksa, R. White III, F. Krueger, A. Kundaje, N. Banovich, and X. Zhang for assistance. We would also like to thank J. Pritchard, C.S. Foo, M. Jones, and other members of the Fraser, Kobor, Koller, and Kundaje Labs for useful discussions and feedback. IMK is funded by the Department of Energy Computational Science Graduate Fellowship Program of the Office of Science and National Nuclear Security Administration in the Department of Energy under contract DE-FG02-97ER25308, the National Science Foundation Graduate Research Fellowship Program under Grant No. DGE-114747, and the Michael J. Flynn Stanford Graduate Fellowship. HBF is an Alfred P. Sloan Fellow and a Pew Scholar in the Biomedical Sciences. MSK is the Canada Research Chair in Social Epigenetics and a Senior Fellow of the Canadian Institute for Advanced Research. This work was supported by NIH grant 1R21HG005240-01A1. This work used the Extreme Science and Engineering Discovery Environment (XSEDE), through the San Diego Supercomputer Center Gordon Compute Cluster, which is supported by National Science Foundation grant number TG-MCB140156 (Towns et al. 2014).

